# Whole-genome sequencing and analysis of two azaleas, *Rhododendron ripense* and *Rhododendron kiyosumense*

**DOI:** 10.1101/2021.06.17.448907

**Authors:** Kenta Shirasawa, Nobuo Kobayashi, Akira Nakatsuka, Hideya Ohta, Sachiko Isobe

**Affiliations:** Kazusa DNA Research Institute, 2-6-7 Kazusa-Kamatari, Kisarazu, Chiba 292-0818, Japan; Faculty of Life and Environmental Science, Shimane University, 1060 Nishikawatsu, Matsue, Shimane 690-8504, Japan

**Keywords:** Chromosome-scale genome assembly, genetic map, long-read sequencing technology, *Rhododendron* subgenus *Tsutsusi* section *Tsutsusi*

## Abstract

To enhance the genomics and genetics of azalea, the whole-genome sequences of two species of *Rhododendron* were determined and analyzed in this study: *Rhododendron ripense*, the cytoplasmic donor and ancestral species of large-flowered and evergreen azalea cultivars, respectively; and *Rhododendron kiyosumense*, a native of Chiba prefecture (Japan) seldomly bred and cultivated. A chromosome-level genome sequence assembly of *R. ripense* was constructed by single-molecule real-time (SMRT) sequencing and genetic mapping, while the genome sequence of *R. kiyosumense* was assembled using the single-tube long fragment read (stLFR) sequencing technology. The *R. ripense* genome assembly contained 319 contigs (506.7 Mb; N50 length: 2.5 Mb) and was assigned to the genetic map to establish 13 pseudomolecule sequences. On the other hand, the genome of *R. kiyosumense* was assembled into 32,308 contigs (601.9 Mb; N50 length: 245.7 kb). A total of 34,606 genes were predicted in the *R. ripense* genome, while 35,785 flower and 48,041 leaf transcript isoforms were identified in *R. kiyosumense* through Iso-Seq analysis. Overall, the genome sequence information generated in this study enhances our understanding of genome evolution in the Ericales and reveals the phylogenetic relationship of closely-related species. This information will also facilitate the development of phenotypically attractive azalea cultivars.

## 1. Introduction

Azalea is a popular woody ornamental plant grown all around the world. In Japan, evergreen azalea varieties (genus *Rhododendron*; subgenus *Tsutsusi*; section *Tsutsusi*), with high ornamental value, were collected from the natural habitat and have served as founding genotypes for breeding phenotypically attractive cultivars since the 17^th^ century^1^. Many of the derived cultivars were introduced into Western countries as new Oriental ornamentals and have been utilized as breeding materials for potted and garden azalea^2^. To determine the phylogenetic relationship among *Rhododendron* and to facilitate the breeding of new varieties with more attractive phenotypes, genome sequence data of the species is needed. However, while the genomic resources of vegetable and fruit crops as well as cereals have dramatically increased because of significant advancements in the next-generation (or 2^nd^ generation) sequencing technologies, following the 1^st^ generation Sanger method^3^, those of woody ornamentals remain limited.

Long-read sequencing technology, also known as the 3^rd^ generation sequencing technology, has revolutionized the field of genomics by improving the contiguity of *de novo* genome sequence assemblies from the contig- and scaffold-level (generated by the 2^nd^ generation sequencing technologies) to the chromosome- or telomere-to-telomere-level^4^. Single-molecule long-read sequencing technologies generate sequence reads containing more than 10k or 100k nucleotides (PacBio, Oxford Nanopore Technologies [ONT]). However, short reads generated by the 2^nd^ generation sequencing technologies could be assembled together using commercially available methods to form long single DNA molecules known as synthetic long reads (Illumina), linked reads (10X Genomics), and single-tube long fragment reads (stLFRs) (MGI tech).

Chromosome-level genome sequence assemblies of two *Rhododendron, R. williamsianum* Rehder and E. H. Wilson and *R. simsii* Planch., have been published recently^5,6^; the former is a typical rhododendron of the subgenus *Hymenanthes*, while the latter is one of the *Rhododendron* native to China. In addition, a draft genome sequence of *Rhododendron delavayi* (subgenus *Hymenanthes*, subsection *Arborea*) is also available^7^. *R. simsii* is considered to be the main ancestral species of Belgian azaleas; however, the results of amplified fragment length polymorphism (AFLP)^8^ and chloroplast DNA (cpDNA) analyses^1^ indicate that *R. simsii* is more distantly related to azalea cultivars, while *Rhododendron ripense* Makino played a greater role in the development of Belgian pot azalea and large-flowered cultivars. *R. ripense*, endemic to West Japan, is the cytoplasmic donor of the large-flowered cultivars^1^ and one of the ancestral species of evergreen azalea cultivars. On the other hand, *Rhododendron kiyosumense* Makino (subgenus *Tsutsusi*, section *Brachycaryx*), a deciduous azalea endemic to Chiba prefecture (Japan), has been seldomly bred and cultivated. In this study, we determined the whole-genome sequences of *R. ripense* and *R. kiyosumense* by single-molecule real-time (SMRT) sequencing and stLFR technology, respectively, to enhance the phylogenetic analysis of azalea cultivars.

## 2. Materials and methods

### 2.1. Plant materials

Leaf samples of *R. ripense* were collected from trees growing at the natural habitat of the Oninoshitaburui Valley (Okuizumo, Shimane, Japan), while those of *R. kiyosumense* were collected from a tree growing in front of the entrance of Kazusa DNA Research Institute (Kisarazu, Chiba, Japan), respectively. Genomic DNA was extracted from the leaf samples of both species using Genomic-Tip (Qiagen, Hilden, Germany). A mapping population of 136 plants was developed from a four-way cross using paternal and maternal genotypes derived from the *Rhododendron* × *hannoense* ‘Amagi-beni-chōjyu’ and *Rhododendron × pulchrum* ‘Oomurasaki’ cross and the *Rhododendron indicum* ‘Chōjyu-hō’ and *Rhododendron obtusum* ‘Kirin’ cross, respectively.

### 2.2. Whole-genome sequencing and assembly of *R. ripense*

*R. ripense* DNA library preparation and sequencing were performed as described previously^9^. Briefly, a short-read DNA library was constructed using the PCR-free Swift 2S Turbo Flexible DNA Library Kit (Swift Sciences, Ann Arbor, MI, USA), which was then converted into a DNA nanoball sequencing library using the MGI Easy Universal Library Conversion Kit (MGI Tech, Shenzhen, China). The library was sequenced using the DNBSEQ G400 (MGI Tech) instrument to generate 101 bp paired-end reads. After removing low-quality bases (<10 quality value) with PRINSEQ and adaptor sequences (AGATCGGAAGAGC) with fastx_clipper in the FASTX-Toolkit, the genome size of *R. ripense* was estimated with Jellyfish. In parallel, a long-read DNA library of *R. ripense* was constructed using the SMRTbell Express Template Prep Kit 2.0 (PacBio, Menlo Park, CA, USA) and sequenced on the PacBio Sequel system (PacBio). The obtained long reads were assembled with Falcon_Unzip to obtain two haplotype-resolved sequences, primary contigs and haplotigs. Potential sequencing errors in the assembled sequences were corrected with ARROW using the long reads. Haplotype duplications in the primary contigs were removed with Purge_Dups. Organelle genome sequences, identified by sequence similarity searches of the reported plastid (DDBJ accession number: MN182619) and mitochondrial (KF386162) genome sequences of *Rhododendron* using Minimap2, were also deleted. Assembly completeness was evaluated with the embryophyta_odb10 data using Benchmarking Universal Single-Copy Orthologs (BUSCO). The software tools used for data analyses are listed in Supplementary Table S1.

### 2.3. Pseudomolecule sequence construction based on genetic mapping

A genetic map of *Rhododendron* was constructed with single nucleotide polymorphisms (SNPs) identified by double digest restriction-site associated DNA sequencing (ddRAD-Seq) of mapping population derived from the four-way cross. Genomic DNA of each genotype in the population was digested with *Pst*I and *Msp*I restriction endonucleases and ligated with adaptors to construct a DNA sequence library, as described previously^10^. The library was sequenced in paired-end mode using the DNBSEQ G400 (MGI Tech) instrument. Sequence reads were subjected to the ddRAD-Seq pipeline^10^ and mapped onto the genome sequence assembly of *R. ripense* with Bowtie2. High-confidence biallelic SNPs were identified using the mpileup option of SAMtools and filtered using VCFtools with the following conditions: read depth ≥ 5; SNP quality = 999; minor allele frequency ≥ 0.2; proportion of missing data < 50%. A linkage analysis of the SNPs was performed with Lep-Map3 to establish a genetic map. The contig sequences were anchored to the genetic map, and pseudomolecule sequences were established with ALLMAPS. The genome structure of *R. ripense* was compared with those of *R. williamsianum, R. simsii*, and 114 additional plant species^3^ whose genome sequences have been assembled at the chromosome level, with minimap2, and visualized using D-GENIES.

### 2.4. Gene prediction and repeat sequence analysis of the *R. ripense* genome

Protein-coding genes were predicted using the MAKER pipeline, based on the peptide sequences of *R. williamsianum* and *R. simsii*, and expressed sequence tags (ESTs) of *R. ripense* (DDBJ accession numbers: FY995432–FY996693). Short gene sequences (<300 bp) as well as genes predicted with an annotation edit distance (AED) greater than 0.5, which is proposed as a threshold for good annotations in the MAKER protocol, were removed to facilitate the selection of high-confidence genes. Functional annotation of the predicted genes was performed with Hayai-Annotation Plants.

Repetitive sequences in the pseudomolecule sequences were identified with RepeatMasker using repeat sequences registered in Repbase and a *de novo* repeat library built with RepeatModeler. The repeat elements were classified into nine types, in accordance with RepeatMasker: short interspersed nuclear elements (SINEs), long interspersed nuclear elements (LINEs), long terminal repeat (LTR) elements, DNA elements, small RNAs, satellites, simple repeats, low complexity repeats, and unclassified.

### 2.5. Whole-genome sequencing and analysis of *R. kiyosumense*

The whole-genome sequence of *R. kiyosumense* was determined using stLFR technology. An stLFR library of *R. kiyosumense* was constructed using the MGIEasy stLFR Library Prep Kit (MGI Tech) and sequenced in paired-end mode using the DNBSEQ G400 (MGI Tech) instrument. The obtained reads were used to estimate the genome size with Jellyfish, as described above. The sequence reads were assembled into contigs with the stLFRdenovo pipeline (https://github.com/BGI-biotools/stLFRdenovo). Sequences from the organelle genomes were eliminated, as described above. The *R. kiyosumense* contig sequences were mapped onto the *R. ripense* pseudomolecule sequences with minimap2.

### 2.6 Transcriptome analysis of *R. kiyosumense*

Total RNA was extracted from flowers and leaves of *R. kiyosumense* using the Plant Total RNA Purification Mini Kit (Favorgen, Ping-Tung, Taiwan). The isolated total RNA was converted into cDNA with NEBNext Single Cell/Low Input cDNA Synthesis & Amplification Module (New England BioLabs, Ipswich, MA, USA). A sequence library for isoform sequencing (Iso-Seq) was prepared using the Iso-Seq Express Template Preparation Kit (PacBio) and sequenced on the PacBio Sequel system (PacBio). Transcript isoforms for each sample were generated with the Iso-Seq3 pipeline (PacBio) implemented in SMRTlink (PacBio). The transcript sequences were aligned to the pseudomolecule sequences of *R. ripense* with Minmap2.

## 3. Results and data description

### 3.1. De novo assembly of the *R. ripense* genome

To estimate the size of the *R. ripense* genome, short reads (39.9 Gb) were subjected to *k*-mer distribution analysis. The results indicated that the *R. ripense* genome was highly heterozygous, with a haploid genome size of 527.0□Mb (Figure 1). A total of 3.0 million sequence reads (58.8 Gb; 111.6x coverage of the estimated genome size) were obtained from four SMRT cells on the PacBio Sequel system. The sequence reads were assembled into primary contigs and alternative sequences, resolved into primary contigs and haplotigs, and then polished to correct potential sequencing errors. Hence, haplotype duplications in the primary contigs as well as possible organelle genome sequences were removed. The final assembly, named RRI_r1.0, consisted of 506.7 Mb of primary contigs (including 318 sequences with an N50 length of 2.5 Mb) and 437.0 Mb haplotigs (including 1,821 sequences with an N50 length of 435.1 kb).

**Figure 1.**
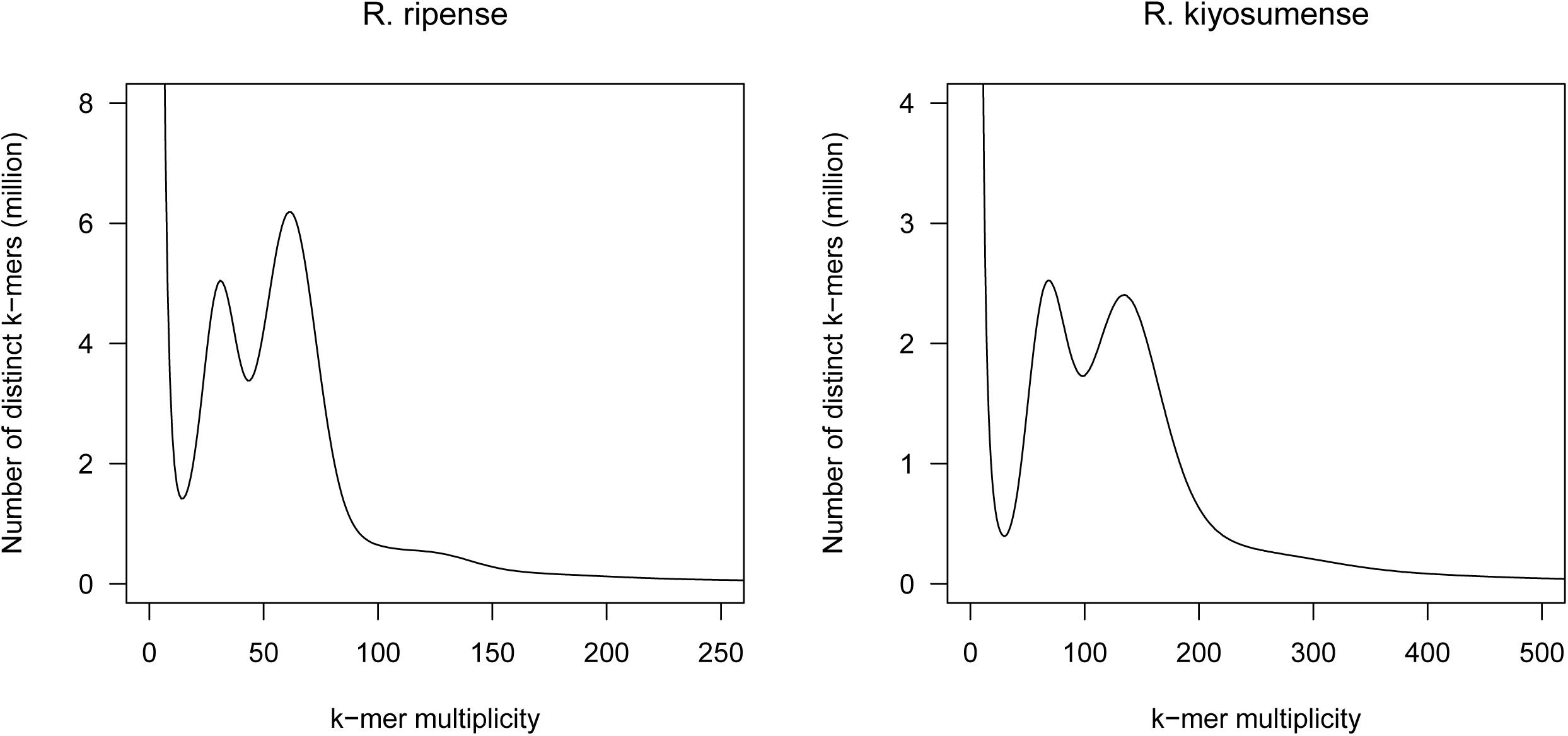
Estimation of the genome size of *Rhododendron ripense* and *Rhododendron kiyosumense*, based on *k*-mer analysis (*k* = □17) with the given multiplicity values.

The genome sequence assembly of *R. ripense* was validated by genetic mapping. To construct a genetic map, 87 million ddRAD-seq reads were obtained from the mapping population (*n* = 136) and parental lines (Supplementary Table S2). High-quality reads were aligned to RRI_r1.0 as a reference, with an average mapping rate of 82.9% (Supplementary Table S2), and 12,463 high-confidence SNPs were detected, which were then employed for linkage analysis. Thirteen linkage groups (LGs), corresponding to the number of chromosomes of *R. ripense*, were obtained, and marker order in each group and map distances between the markers were calculated. The resultant genetic map consisted of 8,723 SNP loci in 1,673 genetic bins spanning a genetic distance of 1,012.3 cM (Figure 2, Table 1, Supplementary Table S3). The nomenclature and orientation of each LG were based on those of *R. williamsianum*^5^. In this mapping process, a probable error in assembly was found in contig Rri1.0p098F_1.1, where upstream and downstream sequences were genetically mapped onto two different linkage groups: LG4 and LG6. Therefore, this contig was split into two sequences, Rri1.1p098F_1.1 and Rri1.1p098F_2.1. The sequence dataset was named RRI_r1.1 (Table 2). The total length of the resultant 319 primary contigs (RRI_r1.1) was 506.7 Mb (96.1% of the estimated genome size) with an N50 length of 2.5 Mb. BUSCO analysis of the primary contigs indicated that 96.9% of the sequences were complete BUSCOs (Table 2). Based on the genetic map, 254 primary contig sequences spanning 487.3 Mb (96.2% of RRI_r1.0, and 92.5% of the estimated genome size) were anchored to the genetic map (Table 3). The sequences were connected with 100 Ns to construct pseudomolecule sequences, namely, the RRI_r1.1.pmol dataset.

**Table 1.** Details of the genetic map of *Rhododendron*.

| Linkage group | Number of SNP loci | Genetic distance<br>(cM) | Genetic bin |
| --- | --- | --- | --- |
| RRI1.1ch01 | 912 | 80.5 | 131 |
| RRI1.1ch02 | 674 | 84.7 | 144 |
| RRI1.1ch03 | 794 | 79.4 | 139 |
| RRI1.1ch04 | 840 | 76.6 | 147 |
| RRI1.1ch05 | 839 | 78.5 | 148 |
| RRI1.1ch06 | 694 | 76.7 | 119 |
| RRI1.1ch07 | 552 | 76.9 | 123 |
| RRI1.1ch08 | 643 | 76.8 | 128 |
| RRI1.1ch09 | 518 | 85.9 | 129 |
| RRI1.1ch10 | 667 | 68.2 | 124 |
| RRI1.1ch11 | 627 | 68.9 | 107 |
| RRI1.1ch12 | 483 | 85.2 | 120 |
| RRI1.1ch13 | 480 | 74.1 | 114 |
| **Total** | **8,723** | **1,012.3** | **1,673** |

**Table 2.** Statistics of the genome assemblies of *R. ripense* (RRI_r1.1) and *R. kiyosumense* (RKI_r1.0).

|  | RRI_r1.1 | RRI_r1.1.haplotigs | RKI_r1.0 |
| --- | --- | --- | --- |
| Total contig size (bp) | 506,708,990 | 437,018,679 | 601,919,567 |
| Number of contigs | 318 | 1,821 | 32,308 |
| Contig N50 length (bp) | 2,455,795 | 435,106 | 245,700 |
| Longest contig size (bp) | 13,760,855 | 4,402,178 | 7,536,965 |
| Gap (bp) | 0 | 0 | 57,426,200 |
| Complete BUSCOs | 96.9 | - | 90.9 |
| Single-copy BUSCOs | 90.3 | - | 83.1 |
| Duplicated BUSCOs | 6.6 | - | 7.8 |
| Fragmented BUSCOs | 0.9 | - | 4.2 |
| Missing BUSCOs | 2.2 | - | 4.9 |
| Number of genes | 34,606 | - | - |

**Table 3.**
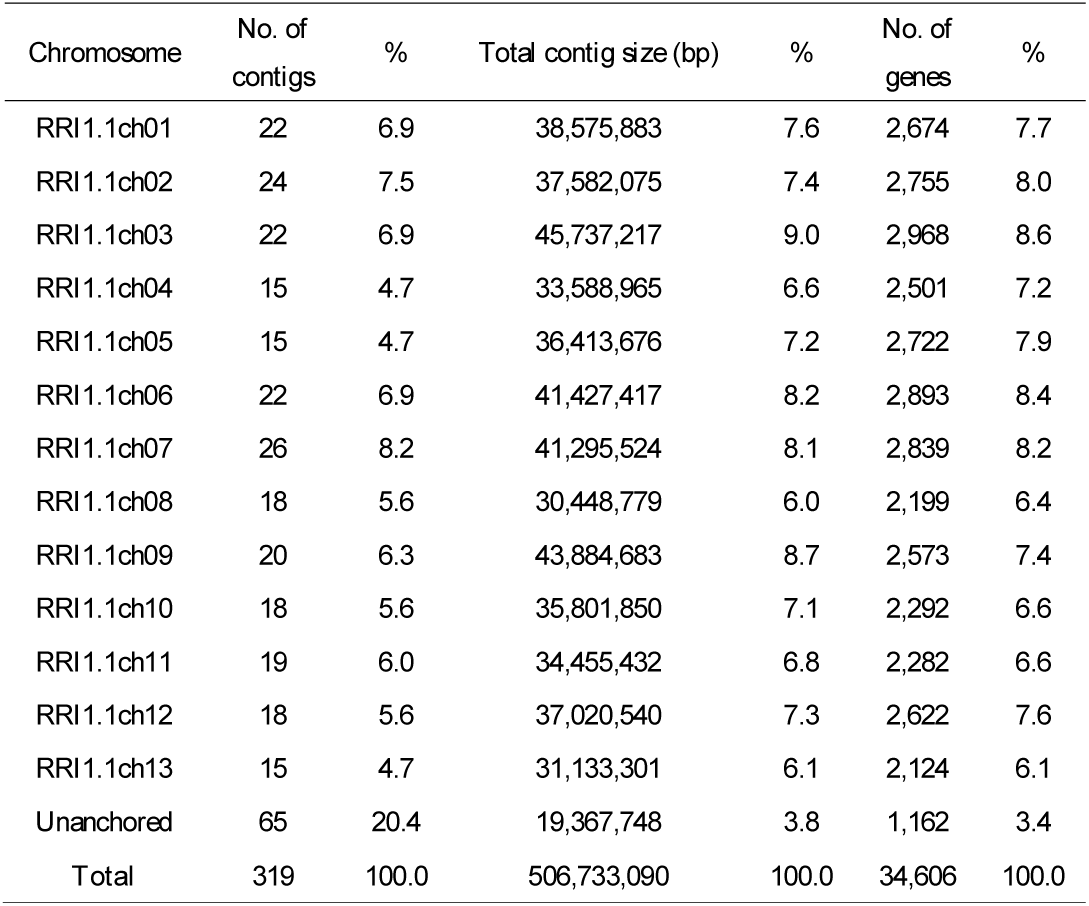
Statistics of the *R. ripense* pseudomolecule sequences.

**Figure 2.**
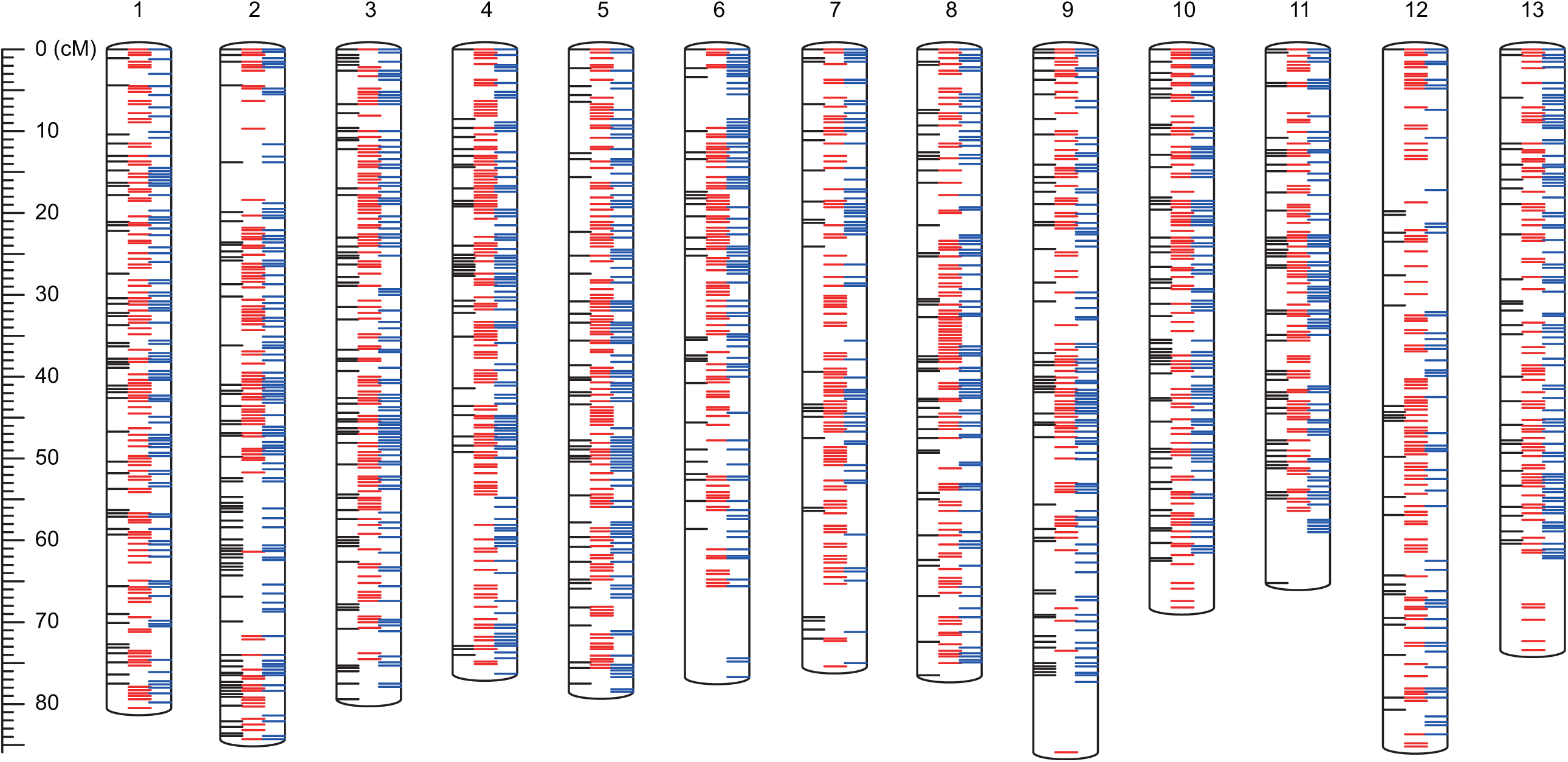
Genetic linkage map of *Rhododendron*. Red and blue bars represent the dominant single nucleotide polymorphisms (SNPs) in the paternal line (F1 hybrid of *R*. × *hannoense* ‘Amagi-beni-chōjyu’ and *R*. × *pulchrum* ‘Oomurasaki’) and maternal line (F1 hybrid of *R. indicum* ‘Chōjyu-hō’ and *R. obtusum* ‘Kirin’), respectively. Black bars indicate co-dominant SNPs between the parental lines. Detailed information of the genetic map and SNP markers is shown in Supplementary Tables S3.

### 3.2. Repetitive sequences and genes in the *R. ripense* genome

Repetitive sequences occupied a physical distance of 260.3 Mb, amounting to 51.4% of the RRI_r1.1.pmol assembly (506.7 Mb). Nine major types of repeats were identified in varying proportions (Table 4). The dominant repeat types in the pseudomolecule sequences were LTR retroelements (77.2 Mb), followed by DNA transposons (38.6 Mb). Repeat sequences unavailable in public databases totaled 116.6 Mb.

**Table 4.** Repetitive sequences in the *R. ripense* genome.

| Repeat type | No. of repetitive elements | Length (bp) | % |
| --- | --- | --- | --- |
| SINEs | 26,849 | 3,854,405 | 0.8 |
| LINEs | 31,674 | 11,213,921 | 2.2 |
| LTR elements | 103,972 | 77,240,248 | 15.2 |
| DNA transposons | 162,900 | 38,589,314 | 7.6 |
| Unclassified | 547,765 | 116,584,817 | 23 |
| Small RNA | 28,312 | 4,355,386 | 0.9 |
| Satellites | 2,068 | 328,499 | 0.1 |
| Simple repeats | 138,314 | 7,082,380 | 1.4 |
| Low complexity | 15,978 | 746,163 | 0.2 |

The initial gene prediction suggested 148,286 putative gene sequences in the *R. ripense* genome assembly. This gene number decreased to 34,606 (Table 2) after the removal of 102,466 low-confidence genes (AED ≥ 0.5), 6,679 genes in repetitive sequence regions, 4,534 short genes (<300 bp), and one redundant sequence. BUSCO analysis of 34,606 genes indicated that 84.6% of the sequences were complete BUSCOs. We concluded that these 34,606 sequences (45.7 Mb in total length) represent high-confidence *R. ripense* genes. Functional annotation analysis of 34,606 genes revealed that 2,313, 3,878, and 2,652 sequences were assigned to Gene Ontology (GO) slim terms in the biological process, cellular component, and molecular function categories, respectively, and 773 genes had enzyme commission (EC) numbers (Supplementary Table S4).

### 3.3. Genome and transcriptome analyses of *R. kiyosumense*

A total of 144.4 Gb reads (100 bp length) were obtained by sequencing the stLFR library of *R. kiyosumense* on two lanes of the MGI DNBSEQ-G400 sequencer. The genome size of *R. kiyosumense* was estimated as 591.6 Mb (Figure 1). The stLFR reads were trimmed to obtain 99.4 Gb clean reads, which were assembled into 32,308 contigs spanning a physical distance of 601.9 Mb, with an N50 length of 245.7 kb (Table 2); the resultant assembly was named RKI_r1.0. The complete BUSCO scores reached 90.9% (Table 2). Of the 32,308 RKI_r1.0 contigs, 22,658 (568.3 Mb) were aligned to the RRI_r1.1.pmol assembly of *R. ripense* (Supplementary Figure S1). Transcript isoforms were obtained in parallel from flowers and leaves of *R. kiyosumense* by Iso-Seq analysis. The obtained Iso-Seq reads were clustered into 35,785 and 48,041 transcript isoforms (mean length: 2,243 bp) in flowers and leaves, respectively. A total of 34,410 flower transcripts and 46,090 leaf transcripts were mapped onto 3,653 and 4,386 contig sequences of RKI_r1.0. Out of the contigs, 391 contigs including 596 flower and 984 leaf isoforms were not assigned to RRI_r1.1.pmol, suggesting that these genome and gene sequences might be absent from the *R. ripense* genome.

### 3.4. Comparative genome analysis of *R. ripense* and its relatives

The genome structure of *R. ripense* was compared with those of *R. williamsianum* and *R. simsii*. In both comparisons, genome structures were well conserved (Figure 3) at the chromosome level, even though the nomenclature and orientation of *R. simsii* chromosomes were different to those of *R. ripense* and *R. williamsianum* chromosomes. The genome structure of *R. ripense* was also compared with those of 114 plant species^3^, whose chromosome-level genome assemblies were publicly available in genome databases. Among these 114 species, *Actinidia chinensis, Actinidia eriantha*, and *Diospyros lotus*, all of which belong to the same order as *Rhododendron* (Ericales), exhibited local synteny along the chromosomes (Figure 3).

**Figure 3.**
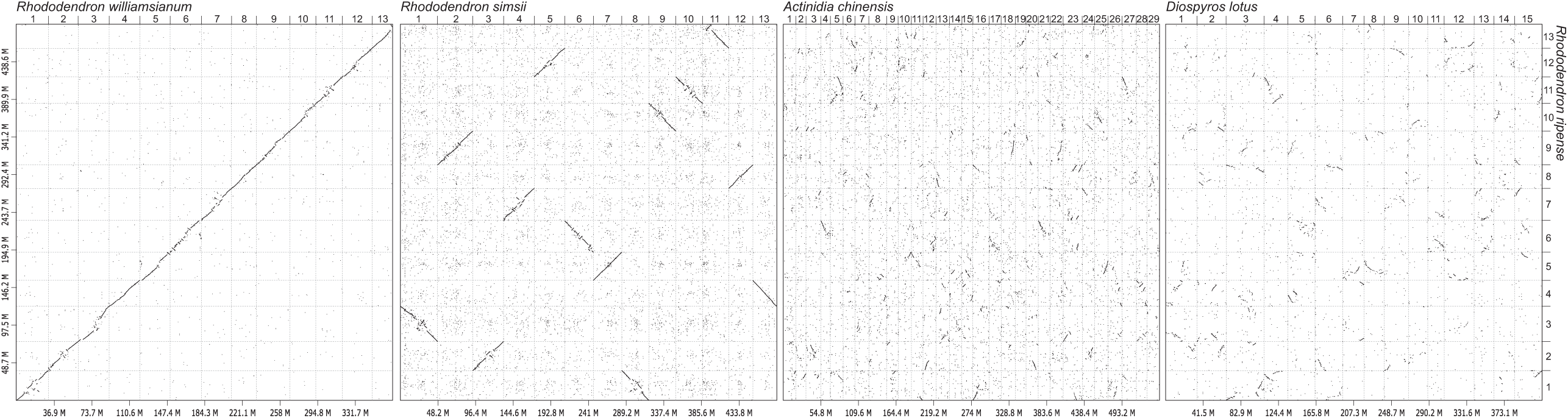
Comparative analysis of the genome sequence and structure of *R. ripense, R. williamsianum, R. simsii*, and 114 additional plant species. Similarities in the genome sequence and structure of *R. ripense, R. williamsianum, R. simsii, Actinidia chinensis*, and *Diospyros lotus* are shown by dots.

## 4. Conclusion and future perspectives

Here, we report the first genome assembly of *R. ripense*, which covered 96.1% of the estimated genome size and included 96.9% of the core gene set of Embryophyta represented by BUSCO (Table 2). The map-based chromosome-level genome sequence covered 96.2% of the assembled sequences (Table 3). This coverage is the highest for any species in the *Rhododendron* (91.1% in *R. simsii* and 68.3% in *R. williamsianum*)^5,6^. In parallel, we established a draft genome sequence of *R. kiyosumense*, with 90.9% complete BUSCOs (Table 2); however, the assembled sequence contiguity represented by N50 was approximately 10 times less than that of *R. ripense* contigs. This difference in contig lengths between the two genomes might be caused by the difference in the sequencing technologies used: the MGI stLFR technique for *R. kiyosumense*, and the PacBio SMRT sequencing technology for *R. ripense*. This possibility is supported by the whole-genome sequencing results of *Macadamia jansenii*, in which the genome assembly generated using the stLFR sequencing technology is more fragmented than that generated by other long-read technologies, e.g., PacBio and ONT^11^. The stLFR could be used to extend genome sequence assemblies with hybrid assemblies using the PacBio and ONT long-read technologies, as suggested by Murigneux et al.^11^

The opposing classification systems proposed by various taxonomists for more than 1,000 species in the *Rhododendron* pose systematic problems at the subgenus level. Recent molecular phylogenetic research, based on several genomic regions, provided a reconstructed phylogeny of the *Rhododendron*^12^. However, the molecular information based on randomly selected genome regions is not sufficient for investigating the phylogenetic relationships of closely-related species in the *Tsutsusi* section. The detailed ancestral relationship between Japanese wild species and traditional cultivars developed by interspecific hybridization remains ambiguous. The genome data generated in this study will serve as a powerful tool to clarify the phylogeny and the breeding history of azaleas. Since bud sport mutants with a different flower color and/or shape represent important azalea cultivars^2^, the genome sequence information could be used to find the causative mutations underlying the attractive trait with high ornament value. The genetic map will also be useful for the identification of several important loci in the *Rhododendron*, which can take more than 3 years (from seeding to flowering) using conventional breeding approaches.

The genome structure of *R. ripense* was conserved not only within the genus *Rhododendron* but also in the order Ericales (Figure 3). The mechanisms of dioecy have been extensively studied in *Actinidia* and *Diospyros*, both of which belong to Ericales^13^. Since *Rhododendron* bear hermaphroditic flowers, comparative genomics in Ericales might provide insights into the molecular mechanisms underlying sex determination in plants. The genome sequence information generated in this study will enhance our understanding of plant genome evolution and will facilitate the development of phenotypically attractive *Rhododendron* cultivars.

## Acknowledgments

We thank Y. Kishida, C. Minami, H. Tsuruoka, and A. Watanabe (Kazusa DNA Research Institute) for providing technical assistance. This work was supported by the Kazusa DNA Research Institute Foundation.

## Data availability

Sequence reads are available from the Sequence Read Archive (DRA) of the DNA Data Bank of Japan (DDBJ) under the accession numbers DRA012151 (whole-genome sequences for *R. ripense*), DRA012152 (ddRAD-Seq for the mapping population), and DRA012155 (whole-genome and transcriptome sequences for *R. kiyosumense*). The DDBJ accession numbers of assembled genome sequences are BPLT01000001-BPLT01000319 (*R. ripense*) and BPLS01000001-BPLS01032308 (*R. kiyosumense*), and those of transcriptome sequences for *R. kiyosumense* are ICRO01000001-ICRO01035785 (flower) and ICRP01000001-ICRP01048041 (leaf). The genome sequence information generated in this study is available at Plant GARDEN (https://plantgarden.jp).

## Supplementary data

**Supplementary Table S1** Software tools used for genome assembly and gene prediction.

**Supplementary Table S2** Double digest restriction-site associated DNA sequencing (ddRAD-Seq) data used for genetic mapping.

**Supplementary Table S3** Genetic map, anchored genome sequences, and single nucleotide polymorphisms (SNPs) in the parental lines of the mapping population.

**Supplementary Table S4** Functional annotations of predicted genes.

**Supplementary Figure S1** Alignment of the *Rhododendron kiyosumense* genome assembly against the *Rhododendron ripense* pseudomolecule sequences. The dot plot shows the collinearity between the genome assembly of *R. kiyosumense* (y-axis) and that of *R. ripense* (x-axis).

