## Supplementary Figure S1 for "Whole-genome sequencing and analysis of two azaleas, *Rhododendron ripense* and *Rhododendron kiyosumense*"

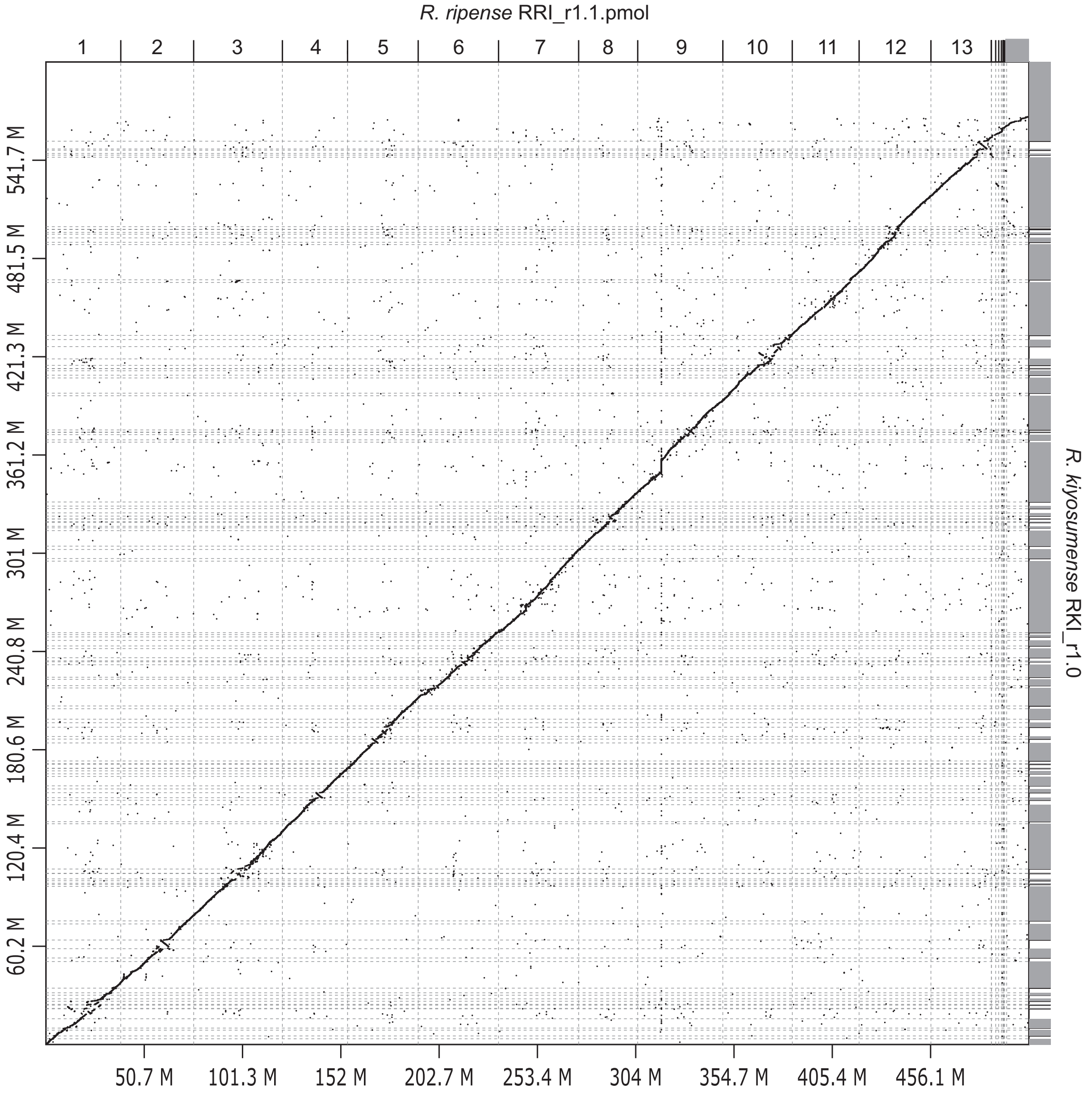


**Supplementary Figure S1** Alignment of the *Rhododendron kiyosumense* genome assembly against the *Rhododendron ripense* pseudomolecule sequences.

The dot plot shows the collinearity between the genome assembly of *R. kiyosumense* (y-axis) and that of *R. ripense* (x-axis).
